# PARG establishes a functional module with BRCA1-BARD1 that controls DNA repair pathway choice during gametogenesis

**DOI:** 10.1101/2022.06.14.496162

**Authors:** Shalini Trivedi, Jitka Blazicková, Nicola Silva

## Abstract

During meiosis, accurate chromosome segregation relies on the formation of programmed DNA double-strand breaks (DSBs). These are in turn repaired by homologous recombination (HR), generating physical attachments between the parental chromosomes called crossovers (COs). Fewer breaks yield recombinant outcomes, while CO-independent mechanisms are employed for repairing the majority of lesions. The balance between different repair pathways is crucial to ensure genome fidelity and to preserve its integrity. We show that *Caenorhabditis elegans* BRC-1/BRCA1-BRD-1/BARD1 and PARG-1/PARG form a complex *in vivo*, that is essential for accurate DNA repair in the germline. Contemporary depletion of BRC-1 and PARG-1 causes synthetic lethality due to reduced CO formation and impaired DSB repair, as evidenced by hindered RPA-1 removal and presence of aberrant chromatin bodies in diakinesis nuclei, whose formation depends on *spo-11* function. These factors largely co-localize and undergo independent loading in developing oocytes, consistent with operating in different pathways. Abrogation of KU- or Theta-mediated end joining elicits opposite effects in *brc-1; parg-1* doubles, highlighting differential involvement of DNA repair pathways and suggesting a profound impact in influencing DNA repair pathway choice by BRC-1-PARG-1. Importantly, lack of PARG-1 catalytic activity suppresses untimely accumulation of RAD-51 foci in *brc-1* mutants but is only partially required to maintain fertility. Altogether, our data show that BRC-1/BRD-1-PARG-1 joined function is essential to keep genome integrity in meiotic cells by regulating multiple DNA repair pathways.

## INTRODUCTION

Genome integrity is constantly challenged by exogenous sources (e.g., chemicals, radiations), however genotoxic insults can also arise within the cellular environment and affect DNA stability, such as upon oxidative stress or errors during replication and transcription.

Severing of both DNA strands (double-strand breaks, DSBs) is one of the most toxic types of DNA damage and if not promptly resolved, it can result in cell death or chromosome aberrations (deletions, translocations), often linked to cellular transformation. Importantly, alterations of the genetic material occurring in the germline can give rise to inheritable mutations, leading to predisposition to cancer and sterility.

During meiosis, DNA replication is followed by two consecutive rounds of cell division, producing haploid gametes that will reconstitute ploidy upon fertilization.

Meiotic progression is characterized by many unique features, all harmonically coordinated to ensure equal partitioning of the genetic material into the daughter cells. The paternal and maternal chromosomes (homologous chromosomes) undergo active motion at meiosis onset until homology is satisfied (homologous pairing), then their association is stabilized by the synaptonemal complex (SC), in whose context homologous recombination can take place and ultimately produce chiasmata (1, 2). These are the cytological manifestation of an occurred crossover (CO), whereby a physical exchange of DNA molecules between the homologs has been achieved.

COs are crucial for the correct segregation of the homologous chromosomes during the first meiotic division, as they act as a physical tether between the homologs that provides the required tension when the pulling forces exerted by the spindles promote the migration of each homolog to the opposite poles of the dividing cell. Failure in CO establishment results in achiasmatic chromosomes that undergo random segregation in the daughter cells, thus causing formation of aneuploid gametes.

COs are formed via HR upon the deliberate induction of DSBs triggered by the topoisomerase-like Spo11 (3), which produces many cuts onto the DNA in order to ensure that at least one for each chromosome pair will be converted into a chiasma. Resolution of the remaining DSBs is channelled into CO-independent, homology-mediated DNA repair pathways (non-CO, NCO) thus restoring genome integrity.

Meiotic DNA repair must be strongly regulated in order to preserve fidelity of the genetic information and therefore it relies on conservative DNA repair mechanisms. Error-prone pathways such as canonical or alternative end joining are normally shut down during meiotic progression, however they can be activated under dysfunctional homologous recombination (4–7).

The tumour suppressor E3 ubiquitin ligase complex BRCA1-BARD1 has been shown to play crucial roles during HR in mitotic cells and in more recent year its roles during meiosis have become increasingly more investigated (8).

In *C. elegans*, BRC-1/BRCA1 and BRD-1/BARD1 are essential to promote inter-sister repair pathway of meiotic DSBs, to stabilize the loading of the recombinase RAD-51/RecA during meiotic progression and recent work has highlighted roles in differentially regulating the recombination rate in oocytes and sperms (8–10).

We have previously shown that the *C. elegans* ortholog of mammalian poly(ADP-ribose)glycohydrolase *PARG, parg-1*, plays crucial roles in ensuring efficient formation of meiotic DSBs as well as promoting their repair via HR during meiosis (11). However, the pathway(s) intersected by PARG-1 function during DSB repair in the germ cells have remained largely unknown.

In a mass spectrometry analysis aimed at identifying novel physical interactors of PARG-1 (12), we have identified BRD-1 as a putative interactor, which forms a obligate heterodimeric complex with BRC-1.

Here, we confirmed that both BRC-1 and BRD-1 co-immunoprecipitate with PARG-1 *in vivo* and contemporary depletion of these factors leads to a severe impairment of fertility and weakening of CO homeostasis, most notably on the X chromosome.

SPO-11-dependent DSBs are formed in *brc-1; parg-1* double mutants however their processing is severely compromised, as indicated by accumulation of RAD-51 foci at late pachytene stage and impaired RPA-1 removal, culminating into generation of morphologically aberrant chromosomes in the diakinesis nuclei whose formation requires SPO-11 function.

We found that abrogation of canonical or alternative non-homologous end joining (cNHEJ and aNHEJ respectively) causes opposite phenotypes in the *brc-1; parg-1* doubles, unveiling an essential function for aNHEJ in preserving fertility levels in both *brc-1* and *brc-1; parg-1* mutants. Furthermore, PARG-1 catalytic activity is responsible for delayed RAD-51 loading in *brc-1* mutants but only partially required to prevent synthetic sterility, indicating that loading of PARG-1 along the chromosomes holds more essential roles in meiotic cells than its enzymatic activity. Our data show that BRC-1-BRD-1 and PARG-1 establish a functional hub that acts as a switch in the DNA repair pathway choice and therefore their joined function is essential to maintain genome integrity during gametogenesis.

## MATERIAL AND METHODS

### Genetics and screenings

*C. elegans* N2 was used as wild type control. Worms were grown on NGM plates according to standard procedures (13) and maintained at 20°C for all the experiments. A full list of the strains used in this study is provided as Supplementary Table 1. Viability and male progeny assessment was performed on single worms of the indicated genotypes, that were transferred onto a fresh plate every 24h. Dead eggs were scored 24h after the mother had been removed and male progeny was counted three days later.

### Immunostaining and image acquisition

Samples were dissected and fixed as in (9) without modifications, except that incubation of primary antibodies was carried out overnight at room temperature instead of 4°C. Primary antibodies used in cytological analyses were: mouse anti-GFP 1:500 (Roche, #11814460001), mouse anti-HA 1:600 (Biolegend, #901513), rabbit anti-HA pre-absorbed against untagged animals 1:100 (Sigma, #H6908), rabbit anti-OLLAS tag pre-absorbed against untagged animals 1:150 (Genscript, #A01658), mouse anti-FLAG pre-absorbed against untagged animals 1:400 (Sigma,, #F1804), rabbit anti-SYP-1 (11) (1:1000), chicken anti-SYP-1 1:400 (15), rabbit anti-BRD-1 pre-absorbed against *brd-1(dw1)* null worms, 1:400 (14), rabbit anti-RAD-51 1:3000 (12), guinea pig anti-HTP-3 1:750 (Y. Kim lab), rabbit anti-HIM-8 1:1000 (Novus Biologicals, # 41980002). All secondary antibodies were Alexa Fluor conjugates to 488, 594 or 647 (ThermoFisher) and were used at 1:500.

Slides were imaged with an upright Zeiss AxioImager.Z2 with 100x objective, equipped with a Monochromatic camera Hamamatsu ORCA Fusion (sCMOS sensor, 2304 × 2304 pixels, 6.5 × 6.5 μm size). Z-stacks were set at 0.24 mm thickness and images were deconvolved using Zen Blue with the “constrained iteration” method set at maximum strength. Images were further processed in Photoshop, where some false coloring was applied. For images in Fig.3C and Fig. S5, samples were acquired with a Delta Vision microscope with a 100x objective, equipped with an Evolve 512 EMCCD Camera and deconvolved with softworx.

For quantification of RAD-51, at least three germlines of the indicated genotypes were used, which were divided into seven equal regions from the mitotic tip to the late pachytene stage. The number of RAD-51 foci was counted in each nucleus and the average was plotted in the charts. The nuclei in late pachytene stage displaying obvious apoptotic features such as chromatin over condensation or reduced nucleus diameter (pyknotic cells), were discarded from the counts. The number of the nuclei quantified in each genotype are reported in Supplementary Table 2.

For synapsis quantification, at least three germlines were used, which were divided into seven equal regions from the mitotic tip to the late pachytene stage. Nuclei were considered fully synapsed only if HTP-3 signal fully overlapped with SYP-1. The number of the nuclei quantified in each genotype are reported in Supplementary Table 2.

For OLLAS::COSA-1 quantification, only nuclei within the last seven rows of cells before diplotene entry were scored. The number of the nuclei quantified in each genotype is reported in Supplementary Table 3.

### DAPI bodies analyses

Worms of the indicated genotypes were selected as L4s and dissected at the indicated times. Animals were dissected and fixed as for regular immunostaining protocol. Once the slides were removed from methanol, they were washed three times in 1x PBST containing 0.1% Tween and 60 µl of a 2µg/ml solution of 4′,6-diamidino-2-phenylindole (DAPI) in water was applied to each slide and left for 1 minute in the dark. Samples were washed for 20 minutes in the dark in 1x PBST and mounted with Vectashield.

For the analysis of the DAPI bodies, only the -1 and -2 diakinesis were included in the counts. The number of the nuclei quantified in each genotype are reported in Supplementary Table 3.

### Immmunoprecipitation and Western blot

Immunoprecipitations were performed by employing nuclear protein extracts containing both nuclear soluble and chromatin-bound fractions produced as in (15). 1 mg of protein extracts was incubated with agarose GFP traps (Chromotek, #GTA-20) pre-equilibrated with Buffer D (20mM HEPES pH 7.9, 150mM KCl, 20% glycerol, 0.2mM EDTA, 0.2% Triton X-100 and complete Roche inhibitor) and carried out on a rotating wheel over night at 4°C. The following day, the samples were spun down at 7500 rpm for 2 minutes and supernatants were discarded. Beads were extensively washed with Buffer D and then resuspended in 40 µl of 2x Laemmli buffer before being boiled for 10 minutes. Eluates were separated from the beads and loaded onto a precast 4-20% acrylamide gel (Bio-Rad), run in Tris-Glycine SDS buffer, and transferred onto a nitrocellulose membrane at 100 V for 90 minutes at 4°C. Blots were blocked in 5% milk in 1x TBS containing 0.1% Tween for 1h at room temperature and primary antibodies were incubated overnight at 4°C. All secondary antibodies were diluted in 5% milk in 1x TBS containing 0.1% Tween and allowed to incubate for 2h at room temperature. Primary antibodies used for Western blot were: mouse anti-HA 1:1000 (Cell Signaling, #2367), mouse anti-PARG-1 1:500 (11), chicken anti-GFP 1:5000 (Abcam, #ab13970), mouse anti-actin 1:1000 (Santa Cruz, #sc-8432), mouse anti-tubulin 1:2000 (Sigma, #T5168).

### Fluorescence *In Situ* Hybridization

FISH was performed as in (15). Briefly, synchronized young adults (24h post-L4) were dissected in 1x EGG buffer (118 mM NaCl, 48 mM KCl, 2 mM CaCl_2_, 2 mM MgCl_2_, 25 mM Hepes, pH 7.3) containing 0.1% tween and fixed with an equal amount of 7.4% of formaldehyde for 2 minutes at room temperature. Coverslips were freeze-cracked in liquid nitrogen and slides were placed in methanol at −20°C for at least 5 minutes. Slides were placed in 50% methanol in 1x SSC for 1 minute at room temperature and then three washes in 2x SSC with 0.1% tween for 5 minutes each were carried out. Samples were placed in 70%, 90% and 100% ethanol at room temperature for 3 minutes each, after which they were air dried. Probes were applied in FISH-mix (10% dextran sulphate, 50% formamide, 2x SSC with 0.1% tween) and hybridization was performed in a slide thermomixer at 93°C for 3 minutes, 72°C for 2 minutes and 37°C overnight. The following day, a 37°C pre-warmed solution of 50% formamide in 2xSCC with 0.1% tween was used to carry two post-hybridization washes of 30 minutes each and then slides were further washed in 2xSCC with 0.1% tween three times for 10 minutes each. Samples were blocked for 30 minutes in a solution of 1% BSA in 2x SSC with 0.1% tween at room temperature and then rhodamine-conjugated anti-digoxigenin (Roche, #11207750910) or FITC-conjugated anti-biotin (AbCam, #ab6650) at a 1:200 and 1:500 dilution respectively, were left to incubate for 3 hours at room temperature in the dark. Slides were washed three times for 10 minutes each in 2xSSC with 0.1% tween in the dark and 60µl of a 2µg/ml solution of DAPI was applied for 2 minutes. Samples were further washed for 20 minutes in 2xSSC with 0.1% tween in the dark and then sealed with Vectashield.

Probes for detection of Chromosome III and Chromosome V were produced from the T17A3 cosmid and by amplifying the 5s rDNA locus by PCR from genomic DNA extracted from N2 wild type animals respectively. The T17A3 cosmid was grown in bacteria and purified with the miniprep kit from Qiagen. Multiple PCR reactions for the 5s rDNA amplicon were pooled and column-purified (Macherey-Nagel). 1µg of DNA was labeled with digoxigenin (for chromosome III) or biotin (for chromosome V) by nick-translation according to manufacturer instructions (Roche, 11745816910 and 11745824910).

### CRISPR/Cas9 genome editing

All transgenic lines were generated by CRISPR as in Paix et al. (16). Briefly, synthetic Ultramers carrying the desired changes were purchased (IDT) and included in a mix also containing tracer RNA and Cas9 protein, the sgRNAs targeting the locus of interest and the *dpy-10* gene (used as a co-editing marker). Tubes were incubated at 37°C for 15 minutes, spun down at 14.000 rpm for 2 minutes and then directly used for injections. WT worms were microinjected (P0) and F1 L4 animals with a *roller* phenotype were individually picked and then used for PCR analysis to identify the heterozygous edits. Non-rollers worms from their progeny were individually picked and PCR were performed to identify the homozygous animals. All strains generated by CRISPR were outcrossed at least twice to N2 wild type animals before use.

## RESULTS

### PARG-1 interacts with BRC-1 and BRD-1 in vivo

We have recently shown that in *C. elegans*, PARG-1/PARG exerts important roles during induction and repair of meiotic DSBs, as removal of *parg-1* from mutant backgrounds with reduced DSBs exacerbates CO defects, while its absence partially alleviates chromosome abnormalities observed in diakinesis nuclei of mutants with impaired HR-mediated repair (11). This indicates that PARG-1 can, either directly or indirectly, not only stimulate DSB formation but also influence DNA repair pathway choice.

In order to shed more light on the repair pathways intersected by *parg-1* function, we took advantage of a mass spectrometry analysis preformed to identify interactors of PARG-1 in meiotic cells, by employing a previously generated functional *parg-1::GFP* tagged line (11). Amongst the putative factors identified, we found several components of the synaptonemal complex and the chromosome axes for which we have previously observed a physical interaction *in vivo* (11), as well as BRD-1 (12), on which we decided to further investigate. Since BRD-1 operates in an obligate heterodimeric complex with BRC-1 both in mammals and nematodes (9, 17), we generated the *brc-1::HA; parg-1::GFP* and *brd-1::HA; parg-1::GFP* strains to perform co-immunoprecipitations experiments, as we previously generated functional tagged lines for these factors (9, 11).

We observed co-immunoprecipitation of both BRC-1 and BRD-1 with PARG-1, confirming the data obtained by mass spectrometry and showing that these proteins form a complex *in vivo* (Fig. 1A). Detection of BRC-1::HA and BRD-1::HA appeared as multiple specific bands (no detection in the untagged WT background was observed), which may indicate alternative isoforms, cleavage and/or partial degradation products generated with this protein extraction/fractionation method (15). Given their physical interaction, we then wondered whether these factors engaged into loading interdependency or influenced each other expression.

**Figure 1.**
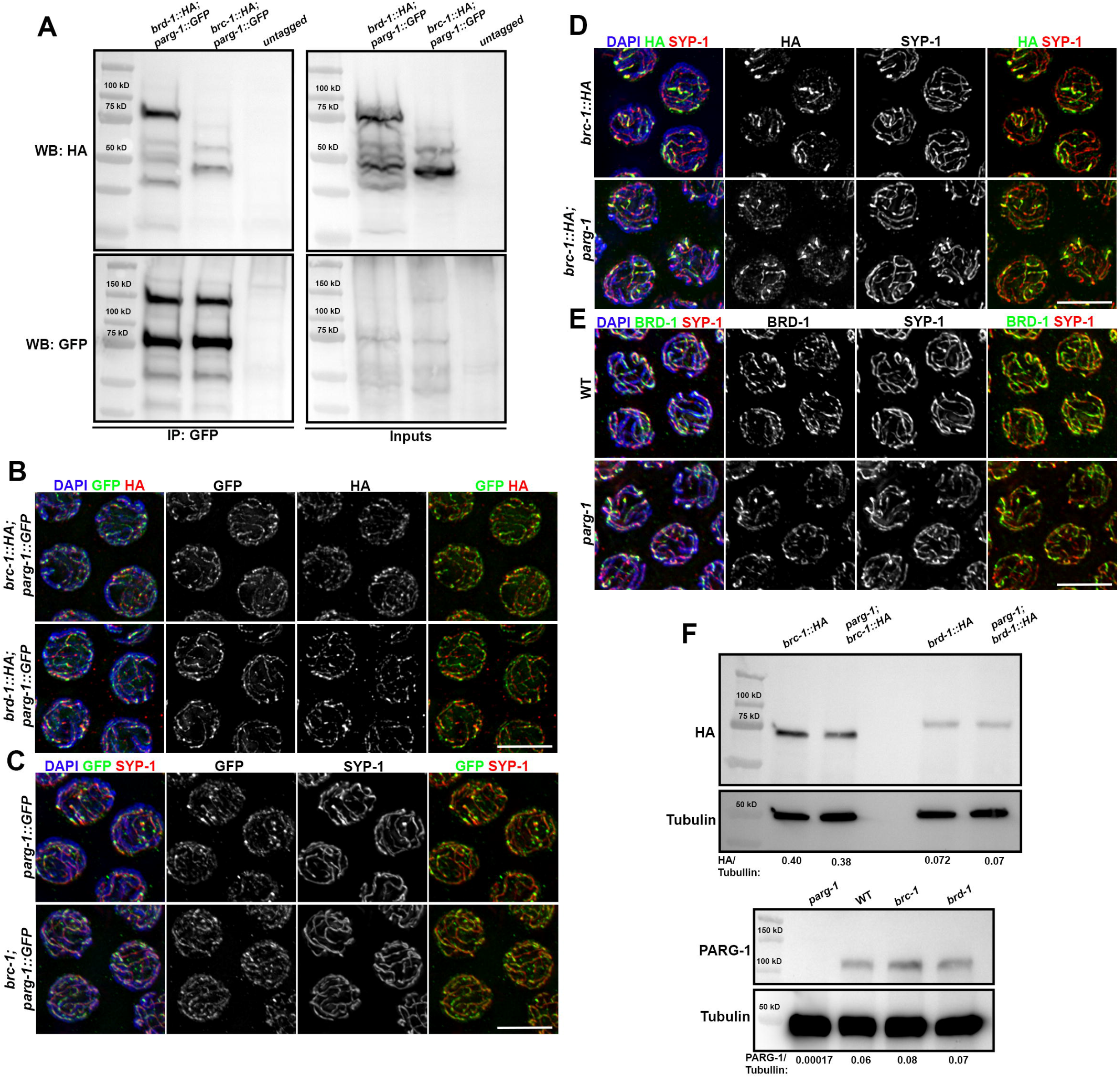
PARG-1 and BRC-1/BRD-1 interact *in vivo*. **(A)** Immunoprecipitation of PARG-1::GFP in brc-1::HA and brd-1::HA strains shows physical interaction. **(B)** Mid-pachytene nuclei stained for HA/GFP showing that BRC-1 and BRD-1 largely co-localize with PARG-1::GFP along synapsed chromosomes. Scale bar 5µm. **(C)** Mid-late pachytene nuclei showing that PARG-1::GFP loading does not require BRC-1. Scale bar 5µm. **(D)** Mid-late pachytene nuclei showing that recruitment of BRC-1::HA and **(E)** BRD-1 along the chromosomes is independent of *parg-1* function. Scale bar 5µm. **(F)** Western blot on total protein extracts showing that absence of *parg-1* does not alter BRC-1 and BRD-1 expression/stability (top), and PARG-1 protein levels are unchanged in *brc-1* and *brd-1* mutants (bottom). Tubulin was used as loading control.

We performed immunofluorescence analysis of PARG-1::GFP and BRC-1::HA/BRD-1::HA (or endogenous BRD-1), which showed that these factors largely co-localize during meiotic Prophase I, switching from a more diffuse nuclear localization in the early stages of meiotic progression to a gradual enrichment along the synapsed chromosomes (9, 11) (Fig. 1B).

Our cytological data indicate that the localization of PARG-1/BRC-1/BRD-1 in the relative mutant backgrounds did not show gross differences when compared to the controls, as PARG-1::GFP was normally loaded in *brc-1(ddr41)* null mutants (Fig. 1C) and conversely, BRC-1::HA (Fig. 1D) or BRD-1 recruitment (Fig. 1E) was not altered in absence of *parg-1*. We extended this analysis by performing Western blot on total worm extracts, which revealed that the stability of BRC-1::HA and BRD-1::HA is not affected in the *parg-1* mutant and PARG-1 levels in *brc-1* and *brd-1* null mutants were similar to wild type controls (Fig. 1F). Thus, we can conclude that PARG-1 engages into a physical interaction with BRC-1-BRD-1 *in vivo*, where these proteins extensively co-localize and are loaded independently of each other.

### Contemporary removal of parg-1 and brc-1 impairs fertility levels

Both *parg-1* and *brc-1* null mutants show nearly normal hatching rates under physiological conditions of growth and consistently, they are fully proficient in homologous pairing, instalment and maintenance of the synaptonemal complex and crossover establishment (9, 11, 18). We then wondered whether removal of both *brc-1* and *parg-1* would elicit synthetic effects during gametogenesis, by generating the *brc-1; parg-1* double mutant strain and assess viability levels.

This analysis revealed high levels of embryonic lethality (Fig. 2A) and segregation of male progeny amongst the survivors (WT, 0/3579 - 0%; *brc-1*, 34/2226 - 1.52%; *parg-1*, 98/3752 - 2.61%; *brc-1; parg-1* 142/1809 - 7.85%) suggesting chromosome segregation defects of both autosomes and the chromosome X respectively (19). While the global embryonic lethality (calculated as the total number of unhatched eggs/total laid eggs) neared 60%, we noticed that the number of dead embryos was not distributed evenly during the typical three days of laying period, but rather showed a steady increase culminating in almost complete sterility at the last day of deposition (day 3, 72hrs), suggesting detrimental age-dependent effects on the fertility levels in the *brc-1; parg-1* background.

**Figure 2.**
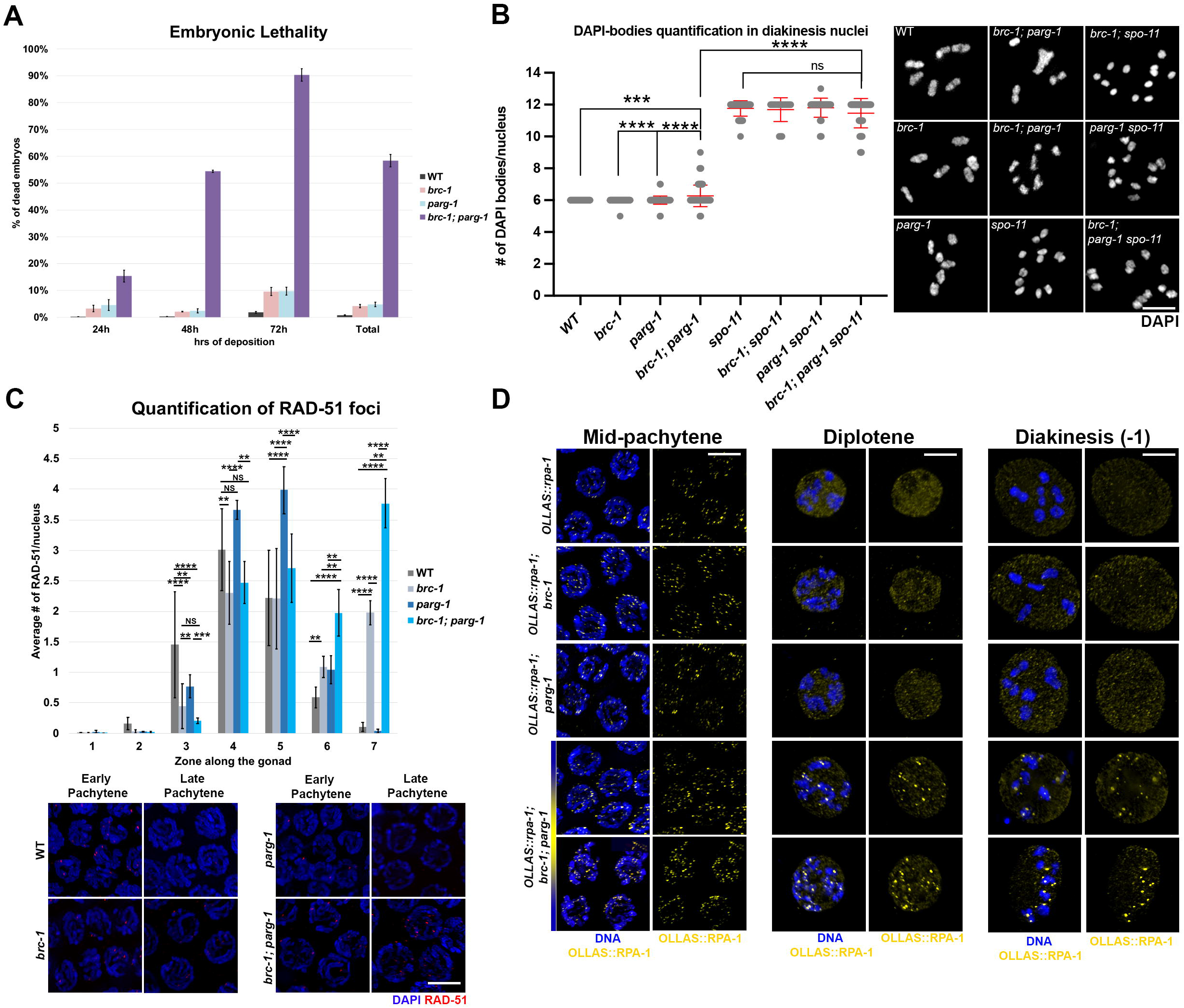
Lack of *brc-1*-*parg-1* causes severe genome instability. **(A)** Assessment of embryonic lethality in the indicated genotypes. The columns in the “total” data point depict sum of all dead embryos/laid eggs over the three-day deposition window. Bars report mean with SEM. **(B)** Quantification of DAPI-bodies (left) and representative images (right) of diakinesis nuclei in the indicated strains. Bars indicate mean with SD, asterisks denote statistically significant differences (*\*\*\*p*=0.0003, *\*\*\*\*p*<0.0001, ^*ns*^non-significant) as calculated by T test. Scale bar 5µm. **(C)** Top: quantification of RAD-51 foci formation in the indicated genotypes. X-axis indicates zone along the gonad, Y-axis shows average number of RAd-51 foci/nucleus. Bars represent mean with SEM and asterisks indicate statistically significant differences (Zone 3:*\*\*p*=0.0036, *\*\*\*\*p*<0.0001, ^*ns*^non-significant; Zone 4:*\*\*p*=0.007, *\*\*\*\*p*<0.0001, ^*ns*^non-significant; Zone 5:*\*\*\*\*p*<0.0001, ^*ns*^non-significant; Zone 6:*\*\*p*=0.002, *\*\*\*\*p*<0.0001, ^*ns*^non-significant; Zone 7:*\*\*p*=0.0059, *\*\*\*\*p*<0.0001, ^*ns*^non-significant) as calculated by Kruskal-Wallis test. Bottom: representative images of nuclei at the indicated stages and genotypes, stained for RAD-51 and DAPI. Scale bar 5µm. **(D)** Nuclei of the indicated genotypes and stages, stained for OLLAS::RPA-1 and DAPI. Scale bar 5µm.

Impairment of different steps during meiotic progression can result in aberrations of chromosome morphology and/or number in the diakinesis nuclei, the last stage of meiotic Prophase I, that can therefore be used as a read out of the events occurred at earlier stages (20). Absent or reduced levels of meiotic DSBs lead to the formation of achiasmatic chromosomes with a regular morphology (univalents) (21–24), which are similarly formed upon i) deficient conversion of the recombination intermediates into post-recombination products (e.g., *msh-4/5* or *cosa-1/CNTD1* mutants) (25–27), ii) or by lack/incomplete synapsis (28, 29).

Dysfunctional DSB repair and deficient end-resection on the other hand, can cause reduced levels of chiasmata which in most instances is coupled with abnormalities of chromosome morphology, including unstructured or fragmented DAPI bodies in the diakinesis nuclei (7, 30–32). Therefore, given the high levels of embryonic lethality observed in the *brc-1; parg-1* double mutants, we sought to perform DAPI-staining analysis of the diakinesis nuclei to assess whether chromosome integrity was affected.

This revealed that lack of both *brc-1* and *parg-1* results in aberrant chromosome morphology and number of DAPI bodies, ranging from unstructured chromatin masses, to univalents and/or fragments (Fig. 2B). Consistent with age-dependent worsening of viability found in the *brc-1; parg-1* double mutants, we observed that the frequency of diakinesis nuclei bearing abnormal chromosome figures was higher in older animals (Fig. S1). Removal of *spo-11* suppressed the formation of morphologically aberrant DAPI-bodies in the diakinesis nuclei of *brc-1; parg-1* doubles (Fig. 2B) leading to formation of normally shaped univalents, suggesting that DSBs are formed but not properly repaired, due to either defective DNA repair pathway/s *per se* or as a consequence of perturbed end resection.

### PARG-1-BRC-1 joined function regulates the processing of recombination intermediates

Given that the analysis of the diakinesis nuclei in *brc-1; parg-1* doubles revealed a highly variable, *spo-11-*dependent phenotype, we then wanted to investigate on both the formation and repair dynamics of recombination intermediates by analysing RAD-51 expression.

RAD-51 is the main recombinase in *C. elegans*, and its loading follows a reproducible pattern in wild type animals. Fewer foci appear at meiotic entry (leptotene/zygotene), they peak at the early-mid pachytene transition and disengage from chromatin at mid-late pachytene stage, indicating completion of repair (28, 33).

It has been previously shown that *brc-1* null mutant hermaphrodites display roughly normal RAD-51 loading in early pachytene but accumulation of foci in late pachytene (9, 10, 18), whereas in *parg-1* mutants RAD-51 accumulates with a slower pace compared to control animals, but peaks above wild type levels in early- and mid-pachytene to then dissipate by late pachytene stage as in wild type worms (11).

Immunostaining of RAD-51 in the *brc-1; parg-1* double mutants mostly recapitulated a similar expression profile as observed in the *brc-1* single mutants except for the late pachytene stage (Fig. 2C, zones 6-7), where a further increase in the number of RAD-51 foci was observed.

To be efficiently loaded in meiotic cells, RAD-51 requires efficient DSB induction (21– 24, 34, 35), presence of single stranded DNA generated upon DSB resection by the MRN/X complex (7, 30), and factors that actively promote RAD-51 loading onto the DNA such as BRCA2/BRC-2 (4, 6).

To assess on whether aberrant RAD-51 loading was caused by perturbations in any of these steps, we analysed localization of i) DSB-1, required for normal DSB induction, ii) MRE-11 and RAD-50, both belonging to the MRN/X complex and iii) BRC-2. We employed available tools for the analysis of MRE-11, while for RAD-50, DSB-1 and BRC-2 we took advantage of functional tagged lines (Fig. S2A-B) purposedly generated for this study by CRISPR/Cas9 approach.

DSB-1, together with DSB-2/-3, has been shown to be essential for DSB induction presumably by promoting a chromatin environment competent for break formation (21, 22, 36). Perturbed DSB induction or recombination, trigger extended expression of DSB-1 until late pachytene, most likely to prolong the “window of opportunity” underlying DSB proficiency. Cytological analysis showed delayed disappearance of 3xHA::DSB-1 in both the *brc-1* and *parg-1* single mutants, as well as in the *brc-1; parg-1* doubles (Fig. S3A), consistent with a repair defect and suggestive of break formation competence (21, 22, 36).

Detection of BRC-2 was previously performed with an antibody directed against the endogenous protein, which showed prominent loading of BRC-2 only upon irradiation (4). Our functional *3xFLAG::brc-2* line however, indicates that BRC-2 is also expressed under physiological conditions of growth at all stages of meiotic prophase I, where it abundantly accumulates in the nucleus (Fig. S3B). We could not detect gross differences in the loading or localization of BRC-2 or MRE-11 and RAD-50 (Fig- S4A-B) neither at early not at later stages of meiotic progression, indicating that the defects in the RAD-51 loading observed in the *brc-1; parg-1* double mutants are likely to stem from a defect downstream DSB induction and resection.

Prior to RAD-51 loading, resected ssDNA is coated with RPA, which binds and stabilizes the DNA filaments before exchanging with RAD-51 (31, 32, 33).

Immunostaining analysis revealed that strikingly, while OLLAS::RPA-1 (40) in *brc-1; parg-1* double mutants was loaded as similarly as in the controls, it did not disengage from DNA at pachytene exit as in WT animals but it rather accumulated in discrete chromatin-associated foci throughout diplotene and diakinesis stages (Fig. 2D). This further corroborates that DSBs are formed and resected in absence of BRC-1 and PARG-1 but their downstream processing is impaired, as presence of chromatin-associated RPA-1 foci indicate lingering ssDNA that did not undergo proper repair.

### BRC-1 and PARG-1 cooperate to ensure robust crossover formation

Given the aberrant processing of the recombination intermediates and increased number of DAPI-bodies in the diakinesis nuclei, we then wondered whether establishment of COs was affected in *brc-1; parg-1* double mutants, by assessing loading of the pro-CO factor COSA-1, which labels the CO designation sites and is essential for their formation (27). Immunofluorescence analysis revealed that ∼60% of the late pachytene nuclei scored displayed the wild type complement of six COSA-1 foci (corresponding to one CO for each homolog pair), whereas the remaining cells bore an aberrant number of foci (Fig. 3A). Most notably, we found that in the remaining 40% of nuclei, the vast majority showed five COSA-1 foci, suggesting lack of a single CO designation site (Fig. 3A).

**Figure 3.**
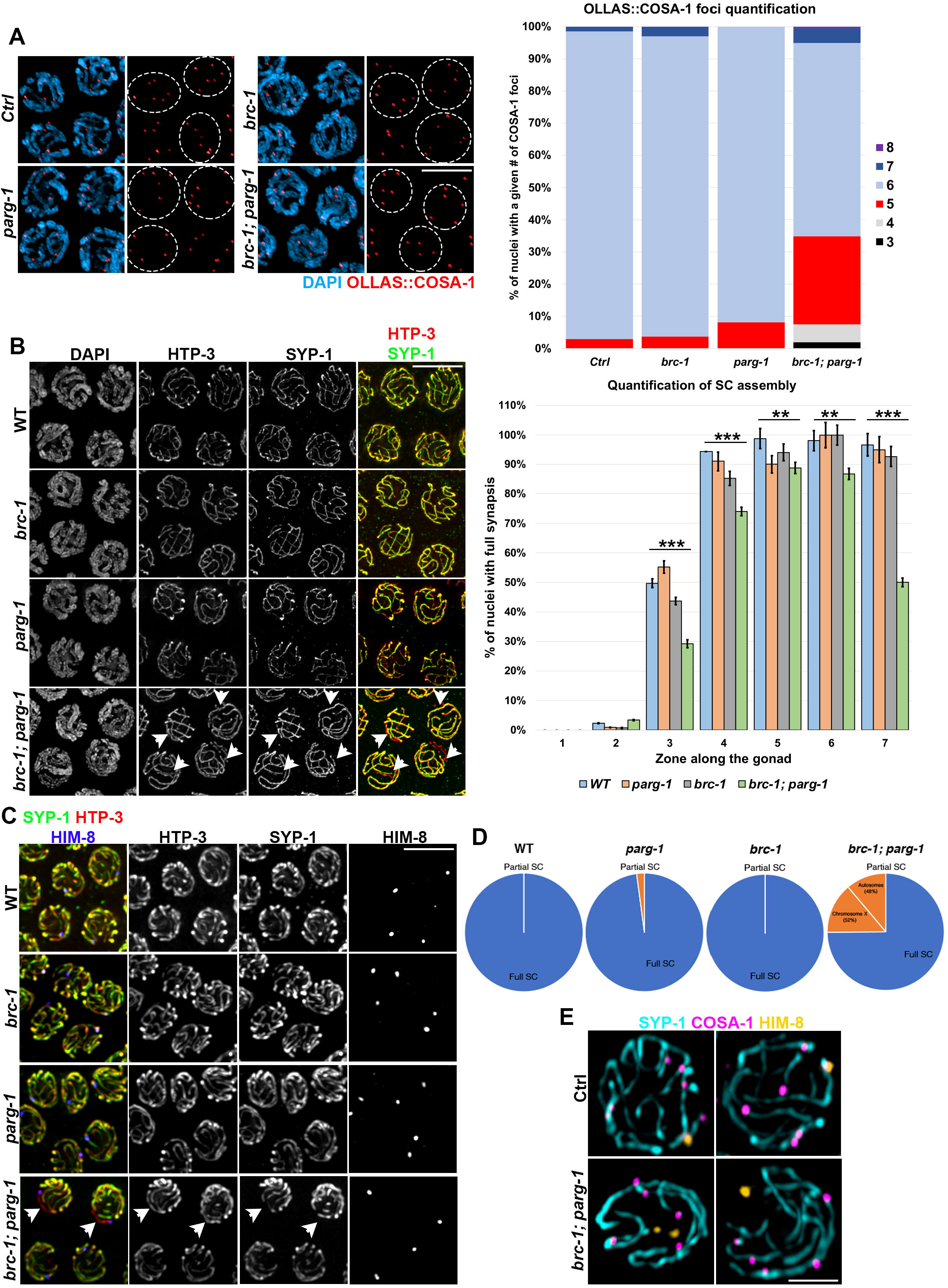
BRC-1-PARG-1 promote robust CO establishment. **(A)** Left: representative images of mid-late pachytene nuclei of the indicated genotypes stained for OLLAS::COSA-1 and DAPI. Dotted circles in the red-channel panels indicate nuclei on the left. Right: quantification of OLLAS::COSA-1 foci number in the last seven rows before diplotene entry. Scale bar 5µm. **(B)** Left: representative images of mid-pachytene nuclei stained for HTP-3/SYP-1 in the indicated genotypes. Arrow heads indicate unsynapsed regions. Right: quantification of SC assembly throughout the gonad in the indicated genotypes. Bars show mean with SEM and asterisks indicate statistically significant differences as calculated by the Х^²^ test (\*\*\**p<0*.*0001*, \*\**p=0*.*0036*). Scale bar 5µm. **(C)** Representative images of mid-pachytene nuclei stained for HTP-3/SYP-1/HIM-8 in the indicated genotypes. Arrow heads indicate unsynapsed chromosome X. Scale bar 5µm. **(D)** Pie charts showing fraction of nuclei with full (blue) or partial (orange) SC. **(E)** Representative images of late pachytene nuclei from controls and brc-1; parg-1 double mutants stained for SYP-1, COSA-1 and HIM-8. Scale bar 5µm.

It has been previously shown that reduced CO numbers trigger the precocious removal of synapsis along the chromosomes that failed in recombining (41, 42), therefore we sought to investigate whether the defective COSA-1 loading was coupled with the untimely removal of the SC in the *brc-1; parg-1* doubles.

To this end, we monitored recruitment of SYP-1 and HTP-3 proteins, localizing to the central elements of the SC and to the chromosome axes respectively (29, 43).

As shown in Fig. 3B, establishment of the SC was slower in *brc-1; parg-1* doubles compared to controls, as well as *brc-1* and *parg-1* single mutants, and upon reaching late pachytene stage (spanning end of zone 6 and overlapping with zone 7), half of the cells analysed displayed chromatin regions stained by HTP-3 but not SYP-1, indicating lack of synapsis. Further, co-staining of the SC and OLLAS::COSA-1 revealed that nuclei displaying de-synapsis carried reduced number of CO designation sites (Fig. S5), confirming that contemporary absence of BRC-1-PARG-1 weakens HR levels.

Next, we sought to investigate on whether the lack of CO and the consequent SC removal were occurring randomly or specifically affecting a chromosome pair.

We monitored the localization of SYP-1, as well as of HIM-8, which binds the sub-telomeric region of the chromosome X (44). If lack of CO was ensuing randomly, we would expect to find de-synapsis of the chromosome X in roughly 1/6 of the cases, since in *C. elegans* there are six pairs of homologous chromosomes. Interestingly, we found that about 50% of the oocytes that displayed early removal of SYP-1 lacked synapsis along the X chromosome (Fig. 3C-D), which consistently displayed no COSA-1 loading as well (Fig. 3E), indicating that the pro-HR activity exerted by the combined action of BRC-1 and PARG-1 is particularly important to establish CO designation sites along the sex chromosomes.

### Abrogation of cNHEJ triggers accumulation of RAD-51 in brc-1; parg-1 mutants and mildly reduces embryonic lethality

During gametogenesis, utilization of potential error-prone pathways for DSB repair is normally prevented and a strong homolog-bias ensures faithful chromosome segregation and maintenance of genome integrity through conservative, “error-free” DNA repair mechanisms such as HR. However, under dysfunctional homologous recombination cells resort to other DNA repair mechanisms to overcome the damage (4, 31, 32). Whether this occurs as an active regulatory process or it is instead due to a constraint imposed by the direct presence of factors operating within the HR pathway, is still unclear (31, 45). Unlike HR, canonical non-homologous end joining (cNHEJ) relies on break-protection and as such, it does not require end-resection. In worms, it proceeds through DSB stabilization operated by the KU70-KU80 heterodimer and Ligase IV-mediated sealing of the breaks (5).

Having established that HR is partially compromised in the *brc-1; parg-1* double mutants, we wanted to address whether cNHEJ played a role in causing genome instability in this background.

To this end, we generated the *brc-1 cku-70; parg-1* triple mutants and analysed RAD-51 staining and diakinesis phenotype.

Previous studies have shown that illegitimate activation of KU-dependent end-joining can cause embryonic lethality and hinder the formation of HR-dependent recombination intermediates during meiotic prophase, leading to reduced numbers of RAD-51 foci that can be restored (at different extents) by abrogation of *cku-70/80* function (4, 6, 31). Assessment of viability levels in the *brc-1 cku-70; parg-1* triple mutants revealed the abrogation of cNHEJ results in a mild, although statistically significant amelioration of embryos viability (Fig. 4A). Interestingly, we found that removal of cNHEJ from both the *brc-1* and *parg-1* single mutants slightly increased embryonic lethality. This suggests that while KU-dependent NHEJ is detrimental under contemporary absence of BRC-1 and PARG-1, it supports, at some degree, fertility levels in the *brc-1* and *parg-1* single mutants (Fig. 4A).

**Figure 4.**
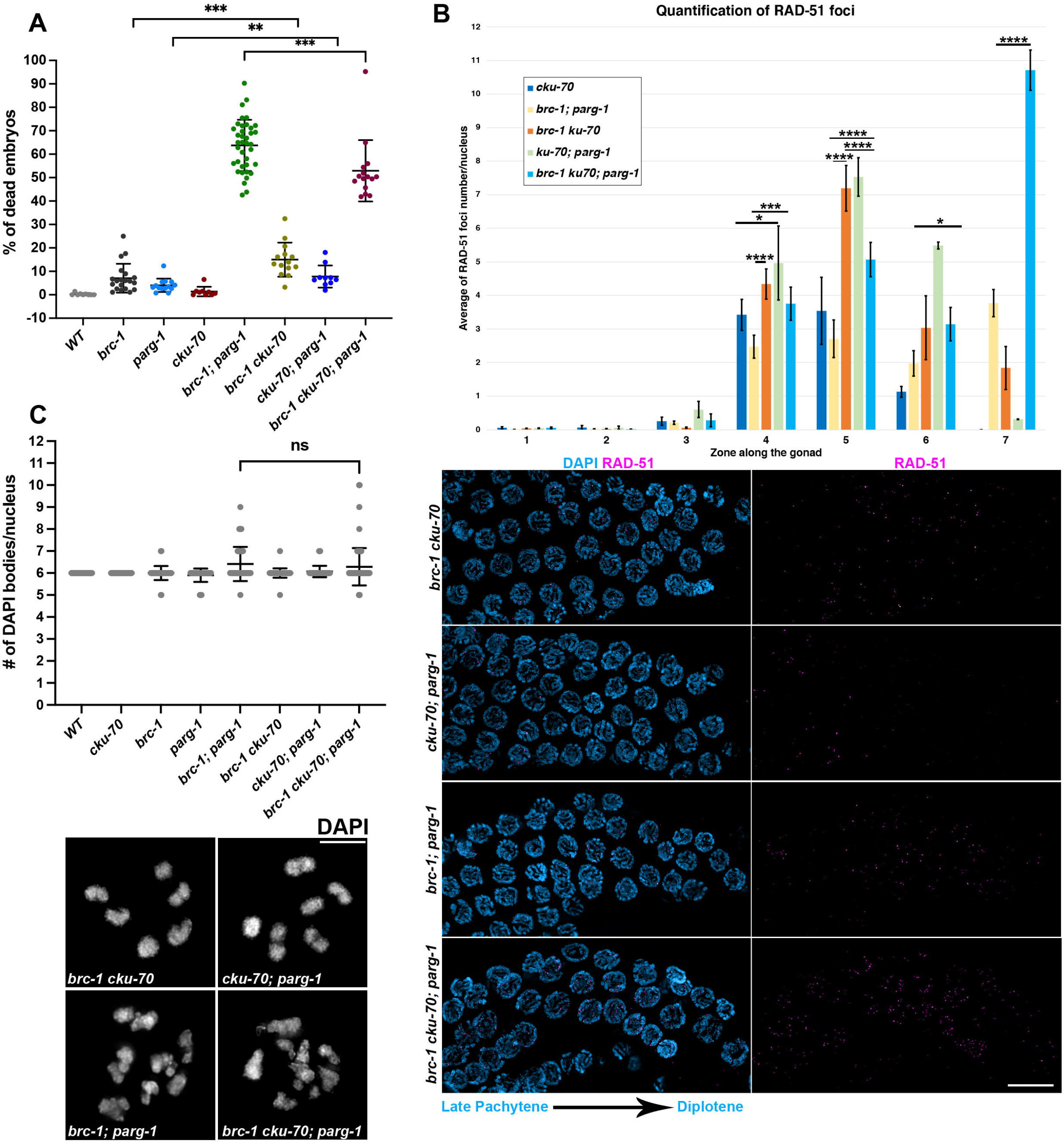
Abrogation of cNHEJ mildly improves viability in *brc-1; parg-1* doubles. **(A)** Quantification of embryonic lethality in the indicated genotypes. Bars indicate mean with SD and asterisks denote statistical significance as calculated by the T test (*\*\*\*p<0*.*0001, **p=0*.*0068)*. **(B)** Top: quantification of RAD-51 foci number in the indicated genotypes across the gonad. Bars show mean with SEM and asterisks denote statistical significance as calculated by the T test (Zone 4: *\*\*\*\*p<0*.*0001, ***p=0*.*0002, *p=0*.*03*; Zone 5: *\*\*\*\*p<0*.*0001*; Zone 6: *\*p=0*.*02*; Zone 7: *\*\*\*\*p<0*.*0001, ***p=0*.*0002, *p=0*.*04*). Bottom: representative images of late pachytene regions from gonads of the indicated genotypes, stained for RAD-51 and DAPI. Scale bar 10µm. **(C)** Top: quantification of DAPI bodies in the indicated genotypes. Bars indicate mean with SD. Bottom: representative images of diakinesis nuclei of the indicated genotypes. Scale bar 5µm.

Analysis of RAD-51 in the *brc-1 cku-70; parg-1* triple mutants revealed a dramatic increase in the number of foci in late pachytene nuclei (Fig. 4B, zone 7), suggesting that removal of cNHEJ redirects a large amount of recombination intermediates into a RAD-51-mediated repair.

However, the increase in RAD-51 loading did not appear to be coupled with a significant improvement of chromatin morphology, since analysis of the diakinesis nuclei in the *brc-1 cku-70; parg-1* triple mutants did not reveal obvious differences compared to the *brc-1; parg-1* doubles (Fig. 4C). Therefore, the fact that aberrant DAPI bodies are still equally formed despite evident engagement of the recombination intermediates into HR-mediated pathway, could suggest either incomplete repair (RAD-51-mediated stranded invasion is inefficient or incomplete) or presence of a significant number of intermediates that cannot be entirely processed via HR in the *brc-1 cku-70; parg-1* triple mutants.

### POLQ-1 exerts essential roles in absence of brc-1 and parg-1

Alternative non-homologous end joining (aNHEJ), also known as TMEJ (Theta-mediated end joining) is independent of KU and requires DNA polymerase Theta/POLQ (*C. elegans polq-1*) (46). TMEJ operates in presence of small regions of homology and consequently it carries an intrinsic mutagenic potential.

In order to address whether TMEJ plays a role in the *brc-1; parg-1* doubles, we built the *brc-1 polq-1; parg-1* triple mutants and performed a similar analysis as for the *brc-1 cku-70; parg-1* mutants.

Removal of *polq-1* resulted in nearly full sterility (Fig. 5A), as the mean of embryonic lethality reached 90% compared to the 60% in *brc-1; parg-1* double mutants. Strikingly, abrogation of *polq-1* functions in *brc-1* mutants caused a synthetic embryonic lethality with a high variability amongst animals, reaching complete lethality in 15% of the animals screened (6/40) and suggesting that in absence of BRC-1, *polq-1* is important to maintain fertility. Removal of *polq-1* increased embryonic lethality also in *parg-1* worms albeit at lesser extent compared to *brc-1* mutant worms, indicating that POLQ-1 exerts some roles in a *parg-1*-depleted background.

**Figure 5.**
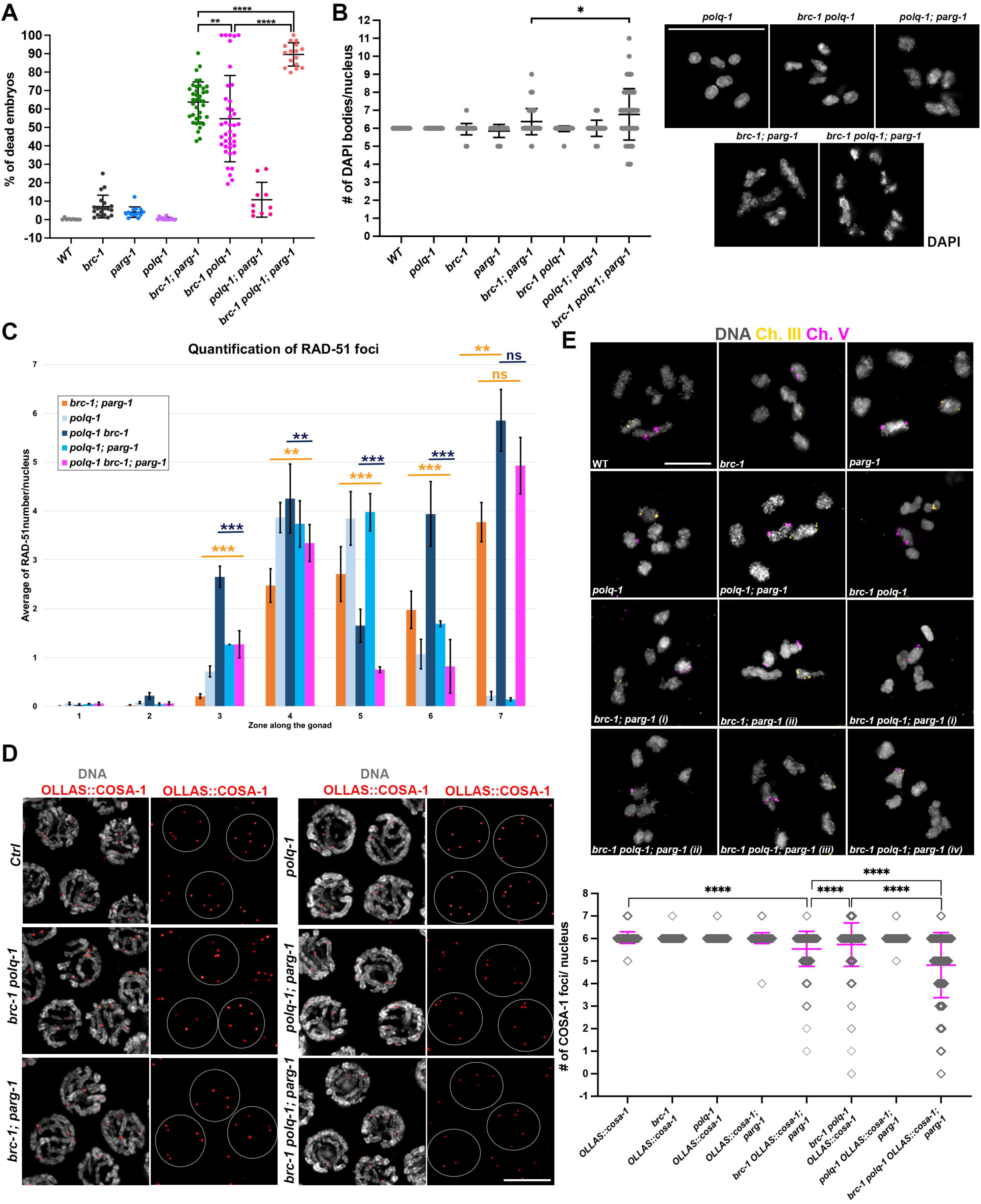
Preventing Theta-mediated function results in synthetic lethality in *brc-1* and *brc-1; parg-1* mutants. **(A)** Quantification of embryonic lethality in the indicated genotypes. Bars indicate mean with SD and asterisks denote statistical significance as calculated by T test (*\*\*\*\*p<0*.*0001, **p=0*.*002*). **(B)** Quantification of DAPI bodies in the indicated genotypes and representative images on the right. Bars show mean with SD and *\*p=0*.*03*. **(C)** Quantification of RAD-51 foci number in the indicated genotypes. Bars show mean with SEM and asterisks indicate statistical significance (Zone 3: ***p<0.0001; Zone 4: *\*\*p=0*.*003*; Zone 5: *\*\*\*p<0*.*0001*; Zone 6: *\*\*\*p<0*.*0001*, Zone 7: *\*\*p=0*.*0032*). **(D)** Representative images of late-pachytene nuclei stained for OLLAS::COSA-1 and DAPI (left) and quantification of OLLAS::COSA-1 foci (right) in the indicated genotypes. Scale bar 5µm. Bars show mean with SD and asterisks denote statistical significance as calculated by the T test (*\*\*\*\*p<0*.*0001*). **(E)** Representative images of FISH analyses on diakinesis nuclei in the indicated genotypes. Scale bar 5µm. Two examples are provided for *brc-1; parg-1* double mutants and four for the *brc-1 polq-1; parg-1* triple mutants.

Analysis of diakinesis nuclei, revealed a significant increase in the number of DAPI bodies in the *brc-1 polq-1; parg-1* triple mutants compared to the *brc-1; parg-1* doubles, indicating that loss of POLQ-1 results in increased chromosome fragmentation or formation of univalents (Fig. 5B). However, chromosome morphology was generally as similarly compromised as observed in the *brc-1; parg-1* double mutants, suggesting that these are not *polq-1*-dependent.

Surprisingly, we did not find superficial aberrations in chromosome morphology or number in the *brc-1 polq-1* worms despite the high levels of embryonic lethality (Fig. 5B).

Assessment of RAD-51 loading revealed that in *brc-1 polq-1; parg-1* mutants there was a significant reduction in the mean number of foci compared to *brc-1; parg-1* doubles in early-mid pachytene (zones 5-6) but not in late pachytene (zone 7). However, at early pachytene stage (zone 4), the average of RAD-51 foci number was comparable between the double and the triple mutants. Furthermore, we noticed that while not having an effect in the *parg-1* mutants, removal of *polq-1* in the *brc-1* mutants triggered higher accumulation of RAD-51 foci compared to *brc-1* single mutants throughout the gonad (Fig. 5C, Fig. 2C).

Given the increased number of DAPI bodies in the diakinesis nuclei, as well as alterations in the RAD-51 loading in the *brc-1 polq-1; parg-1* triple mutants, we wondered whether these phenotypes had an effect also on the CO-designation sites. To this end, we analysed COSA-1 recruitment in the triple mutants, which revealed presence of several nuclei with reduced COSA-1 foci in the *brc-1 polq-1* doubles, although their distribution was not as affected as in the *brc-1; parg-1* mutants (Fig. 5D). Strikingly, the *brc-1 polq-1; parg-1* triple mutants displayed further reduced COSA-1 loading in comparison to *brc-1; parg-1* and *brc-1 polq-1* mutants, indicating that POLQ-1 promotes, most likely indirectly, formation of CO intermediates when BRC-1-PARG-1 function is compromised.

Reduced levels of HR-dependent repair were also further highlighted by formation of non-homologous chromosome fusions observed in the diakinesis nuclei of *brc-1; parg-1* and *brc-1 polq-1; parg-1* mutants through FISH analysis, in which we monitored DAPI-bodies identity by employing specific probes for the autosomes III and V (Fig. 5E).

Furthermore, analysis of SC assembly also revealed that in the *brc-1 polq-1; parg-1* mutants, establishment of synapsis occurs much slower than in the *brc-1; parg-1* doubles but reaches comparable levels in late pachytene (Fig. S6). Interestingly, we found that in the *brc-1 polq-1* double mutants, while the kinetics of the SC assembly were not significantly affected at early stages, significant levels of desynapsis were found in late pachytene as for *brc-1; parg-1*, further indicating a failure in achieving robust CO establishment (Fig. S6) and well aligning with the reduced COSA-1 loading (Fig. 5D).

Taken together, these data unveil a complex, multi-layered control exerted on DNA repair pathway choice by BRC-1 and PARG-1 that intersects both KU- and Theta-mediated end joining pathways, whose functional ramifications differently impact on HR-mediated repair.

### PARG-1 catalytic activity partially safeguards genome integrity in absence of brc-1

We have previously shown that PARG-1 loading along the chromosomes and its catalytic activity in removing poly(ADP)ribose (PAR) moieties hold differential roles during gametogenesis in worms. Loading of PARG-1 is essential to perform its functions in promoting proper DSB induction and HR-mediated repair, whereas PARG-1 catalytic activity, while having an effect in regulating the protein turn over, does not seem to be required for recombination in backgrounds with reduced DSBs (11).

Given the synthetic effects observed in the *brc-1; parg-1* double mutants, we sought to investigate on whether PARG-1 catalytic activity or rather its impaired loading were the underlying cause of the defects that we observed.

To this end, we generated the *brc-1; parg-1(CD)* double mutants and assessed embryonic viability and RAD-51 foci formation.

We found that abrogation of PARG-1 catalytic activity in *brc-1* mutants caused a 3-folds increase in the embryonic lethality, however, the percentage of dead embryos was roughly half compared to *brc-1; parg-1* null worms (Fig. 6A), suggesting that the catalytic activity is only partially required to prevent lethality.

**Figure 6.**
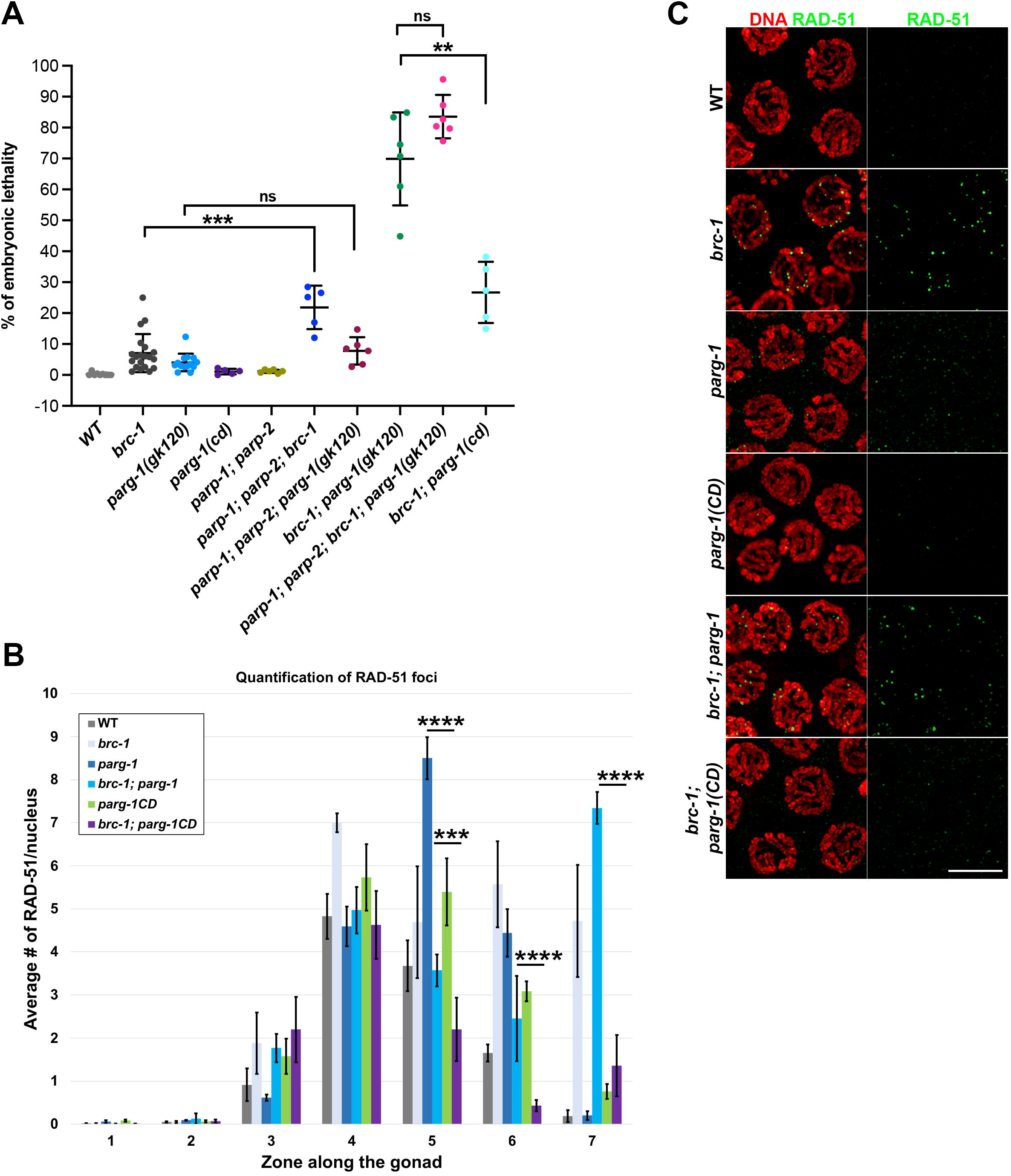
PARG-1 catalytic activity is partially required to avert genome instability in absence of *brc-1*. **(A)** Quantification of embryonic lethality in the indicated genotypes. Bars show mean with SD and asterisks indicate statistical significance as calculated by the T test (*\*\*\*p<0*.*0001, **p=0*.*0002*, ^*ns*^*non-significant*). **(B)** Quantification of RAD-51 foci number and representative images of mid-pachytene nuclei **(C)** in the indicated genotypes. Bars show mean with SEM and asterisks denote statistical significance as assessed by the T test (*\*\*\*\*p<0*.*0001, ***p=0*.*00033*). Scale bar 5µm.e

Next, we generated the *parp-1; parp-2; brc-1; parg-1* quadruple mutants in which PAR synthesis was abolished by abrogating the function of the poly(ADP)ribose polymerases PARP-1 and PARP-2 (11, 47). This did not result in a rescue or attenuation of the embryonic lethality, further corroborating that in absence of BRC-1, obliterating PAR synthesis or preventing its removal has only partial effects on genome stability, and that loading of PARG-1 along chromosomes is likely to hold more crucial functions during meiosis.

Strikingly, analysis of RAD-51 recruitment showed that PARG-1 catalytic activity is essential for the formation of foci at mid-late pachytene stages observed upon BRC-1 removal (Fig. 6B-C), indicating that either the kinetics of the recombination intermediates processing or of their formation is regulated by the enzymatic function of PARG-1.

Taken together, our data show that BRC-1 and PARG-1 joined function has a profound activity in modulating DNA repair pathway choice during oocyte development, exerting crucial roles as safekeepers of genome integrity in the gametes.

## Discussion

ADP-ribosylation is a crucial post-translational modification that plays pivotal roles during DNA repair. The founder “writers” and “erasers”, PARP1/2 and PARG respectively, have been extensively studied in mitotic cell cultures models, however their characterization during gametogenesis has received significantly less attention, since mice mutants of PARG and double knockouts of PARP1/2 are embryonic lethal (48–52), thus hindering their study in the germline.

Inhibition of poly(ADP)-ribosylation (PARylation) in cancer cells harbouring mutations in the BRCA1/2 genes elicits synthetic lethality due to impaired repair of DSBs arising from the collision of the replication forks with trapped PARP1-DNA adducts. This synthetic lethality is currently exploited in cancer therapy (53–56). A similar approach aimed at targeting PARG has proven to be more problematic, due to lower specificity and high toxicity of available chemical inhibitors (56). Therefore, whether impacting on PARG function elicits the same phenotypes observed for PARPs inhibition in BRCAs-mutated cells, is still under debate.

We show that abrogating PARG function in null *brc-1* mutant worms causes high embryonic lethality due to aberrant DNA repair and weakened CO establishment in developing oocytes. Contemporary loss of *brc-1* and *parg-1* results in the formation of aberrant chromosome structures in the diakinesis nuclei, indicating that BRC-1 and PARG-1 are likely to act in different pathways during meiotic progression.

Abrogation of *brc-1* function perturbs RAD-51 loading in the *C. elegans* gonad, where it accumulates in chromatin associated foci at late pachytene stage (10). In contrast, in *parg-1* mutants the number of RAD-51-labeled recombination intermediates rises above WT levels at early-mid pachytene stages (11). RAD-51 dynamics resemble *brc-1*’s at early pachytene in the *brc-1; parg-1* double mutants, indicating that BRC-1 may act upstream PARG-1 in influencing formation/processing of recombination intermediates, and show a synthetic effect in late pachytene.

Reduced RAD-51 foci number in *brc-1; parg-1* versus *parg-1* single mutants is unlikely to originate from impaired DSB formation or reduced end-resection, since we found that loading of the pro-DSB factor DSB-1 and the MRN complex components MRE-11 and RAD-50 respectively, do not display superficial differences compared to controls. Our data indicate that meiotic DSBs are formed in absence of *brc-1* and *parg-1* but are not efficiently repaired, as further corroborated by normal RPA-1 loading but impaired removal, and nonhomologous chromosome fusions observed in diakinesis nuclei. Deletion of SPO-11 in the *brc-1; parg-1* doubles results in the formation of normally shaped univalents, emphasizing that aberrant chromosome figures depend on presence of meiotic DSBs. Moreover, we can rule out that blocked RPA-1 removal is not caused by hindered exchange with RAD-51 *per se*, which would prevent RAD-51 loading altogether.

We envision a scenario whereby blocking BRC-1/PARG-1 function leads to the formation of recombination intermediates that are not a substrate for HR, which in turn results in DSBs being hijacked by multiple illegitimate error-prone pathways. It has been postulated that BRC-1 activity promotes non-crossover-mediated repair and that in its absence, delayed recombination intermediates may be channelled into alternative forms of repair. PARG-1 could be involved in promoting such alternative repair pathway, although it is worth mentioning that a large proportion of nuclei in the *brc-1; parg-1* double mutants display a normal complement of morphologically normal bivalents in the diakinesis nuclei, as well as proper numbers of CO-designation sites, indicating that PARG-1 could act on a subset of recombination intermediates. Intriguingly, we find that lack of COs is particularly prominent on the chromosome X, suggesting that PARG-1 functions in promoting recombination might be required at different extents between autosomes and sex chromosomes. This is consistent with our previous finding that in *him-5* mutants, which are defective in forming a CO on the chromosome X but not on the autosomes, lack of *parg-1* weakens IR-dependent rescue of chiasmata (11).

Preventing KU-mediated non-homologous end joining induces a dramatic formation of RAD-51 foci in late pachytene cells and mildly alleviates embryonic lethality in the *brc-1; parg-1* double mutants. However, this does not seem to be coupled with a significant improvement of chromosome morphology in the diakinesis nuclei. Activation of cNHEJ has been often observed under dysfunctional HR, which under normal conditions could act by directly preventing binding of KU to dsDNA and therefore allowing conservative DNA repair pathways to take place (4, 6, 31). On the other hand, removal of *polq-1* causes high embryonic lethality in *brc-1* mutants and complete sterility in the *brc-1; parg-1* doubles, indicating that in absence of BRC-1, TMEJ is essential to maintain fertility. Surprisingly, we did not find obvious alterations in the morphology or number of DAPI bodies in *brc-1 polq-1* double mutants despite the high level of embryonic lethality. This could indicate that the outcome of the aberrant repair taking place in these double mutants may not be detectable at the level of chromosome morphology in the diakinesis nuclei, and it could be unveiled at later stages of meiotic progression such as in Metaphase I-Anaphase I (e.g., lagging chromosomes, chromatin bridges), preventing proper chromosome segregation. Interestingly, RAD-51 foci were overall increased in number in the *brc-1 polq-1* double mutants, suggesting that removal of POLQ-1 triggers accumulation of recombination intermediates at all stages of Prophase I.

Previous work has shown that lack of *brc-1* triggers accumulation of mutations throughout generations, in a *polq-1*-dependent fashion (57), thus the worsening in the viability levels that we observe, seems rather counterintuitive. However, it is possible that while abrogating *polq-1* function in a *brc-1*-deficient background suppresses its mutagenic potential, at the same time it may also affect genome stability by other means. In fact, the *polq-1*-dependent impact on genomic fidelity could arise by other (*polq-1*-independent) sources, that result in the activation of TMEJ as an emergency resource mechanism. Hence, removing POLQ-1 would avert formation of mutations, but at the same time could also suppress an instrumental tool of repairing DNA damage, ultimately resulting in reduced viability. We find that indeed that the *brc-1 polq-1; parg-1* triple mutants show almost complete embryonic lethality, display incomplete chromosome synapsis and bear a further reduction in COSA-1 loading, suggesting that POLQ-1 is responsible for maintaining residual levels of viability in absence of *brc-1* and *parg-1* and further, to promote establishment of CO-designation sites, although most likely indirectly.

The presence of chromosome aberrations upon removal of *cku-70* or *polq-1*, irrespectively of the effects on RAD-51 loading, indicates that multiple repair pathways are engaged in the *brc-1; parg-1* double mutants and still act despite removal of *cku-70* or *polq-1*, whose activity results in chromatin abnormalities.

This would hint at the possibility that a plethora of pathways can act on meiotic breaks and therefore giving rise to a high redundancy, as also recent work has indeed shown in the *C. elegans* germ line (45).

Interestingly, we found that contrary to what we previously observed in *him-5* mutants (11), PARG-1 catalytic activity plays pivotal roles in promoting RAD-51-labeled recombination intermediates in late pachytene cells in absence of *brc-1*, and this is coupled with increased embryonic lethality. However, in *brc-1; parg-1* null worms the embryonic lethality is much higher, thus we can conclude that while playing important activities in promoting repair of late recombination intermediates generated upon deficient *brc-1* function, yet the enzymatic activity of PARG-1 is only partially required to safeguard genome stability in absence of BRC-1.

In conclusion, our data demonstrate that as observed upon PARPs inhibition in mitotic cells, blocking PARG-1 functions in *brc-1* mutated animals confers synthetic effects. Our work shows that abrogation of PARP-1 and PARP-2 function however, has less dramatic repercussions on *brc-1* mutants than lack of PARG-1. This could be due to the fact that PARPs inhibition and PARPs knockout are likely to elicit very different responses, since in the former instance the proteins remain trapped on the chromatin, whereas in the latter case they are not present in the cells.

In fact, this aligns with the findings in human mitotic cells whereby knock down of *PARP1/2* in *BRCA*^-/-^ cells does not elicit the same extent of lethality as upon PARP1 inhibition, indicating that the mechanisms underlying these responses are very likely to be different.

We have uncovered an intricated and complex activity mediated by BRC-1-PARG-1 in regulating DNA repair pathway choice during gametogenesis, with functional ramifications reaching out to both homologous recombination as well as non-CO dependent repair, placing these two important factors at a crossroad between multiple routes.

## Data Availability

All the relevant data underlying the experiments shown in this study are included in the manuscript. Requests for strains and/or reagents should be addressed to the corresponding author.

## Acknowledgments

We are grateful to S. Boulton for the anti-BRD-1 antibody, E. Martinez-Perez for the chicken anti-SYP-1, A. Villeneuve for the *rad-50(ok197)* mutant and S. Smolikove for the *mre-11::GFP* and *OLLAS::rpa-1* strains. We are grateful to Angela Graf and Verena Jantsch for the unvaluable help in generating the *3xHA::dsb-1, 3xFLAG::brc-2* and *rad-50::3xFLAG* tagged lines. We acknowledge the core facility CELLIM supported by the Czech- BioImaging large RI project (LM2018129 funded by MEYS CR) for their support with obtaining scientific data presented in this paper.

## Author contributions

NS designed the research; ST and NS performed the experiments with the technical support of JB; NS supervised the project and wrote the manuscript.

## Supplementary Figure Legends

**Supp. Fig.1. Chromosome morphology worsens with age in brc-1; parg-1 double mutants**.

Top: quantification of DAPI bodies in the indicated genotypes, in 48h post-L4 worms. Bars indicate mean with SD and asterisks indicate statistical significance as calculated by the T test (*\*\*\*\*p<0*.*0001*). Bottom: representative images of diakinesis nuclei in the indicated genotypes. Scale bar 2µm.

**Supp. Fig.2. Functionality assessment of *rad-50, brc-2* and *dsb-1* tagged lines**.

**(A)** Quantification of embryonic lethality in the indicated genotypes. **(B)** Quantification of embryonic lethality in worms exposed to the indicated IR dose and under untreated conditions. Young adults were exposed or not to IR, allowed to lay eggs for 24h and then mothers were sacrificed and dead eggs counted 24h afterwards.

**Supp. Fig.3. *brc-1* and *parg-1* are dispensable for DSB-1 and BRC-2 localization**.

**(A)** 3xHA::DSB-1 localization is extended in the *brc-1; parg-1* double mutants. Scale bar 20µm. **(B)** *brc-1; parg-1* worms display normal localization of 3xFLAG::BRC-2. Scale bar 20µm.

**Supp. Fig.4. *brc-1* and *parg-1* are dispensable for MRE-11 and RAD-50 localization**.

**(A)** MRE-11::GFP localization does not display aberrancies in the *brc-1; parg-1* double mutants. Scale bar 20µm. **(B)** *brc-1; parg-1* worms display normal localization of RAD-50::3xFLAG. Scale bar 20µm.

**Supp. Fig.5. BRC-1-PARG-1 promote establishment of CO on the chromosome X**.

Mid-late pachytene nuclei stained for SYP-1/HTP-3/COSA-1 in the indicated genotypes. Numbers in parentheses refer to COSA-1 foci. Note that unsynapsed chromosomes lack COSA-1 focus. Scale bar 5µm.

**Supp. Fig.6. Lack of POLQ-1 weakens SC establishment in *brc-1* and *brc-1; parg-1* mutants**.

Left: quantification of SC assembly across the gonad of the indicated genotypes. Bars indicate % of nuclei with full SC and SEM, asterisks denote statistical significance as calculate by the Х^2^ test (*\*\*\*\*p<0*.*0001, Zone 3: **p=0*.*003, Zone 6: p=0*.*002*, ^*NS*^*non-significant*). Right: representative images of mid-late pachytene nuclei stained for HTP-3/SYP-1 in the indicated genotypes. Scale bar 5µm.

