## Supplementary figures and images for "PARG establishes a functional module with BRCA1-BARD1 that controls DNA repair pathway choice during gametogenesis"

### Supplemnetal Figure 1

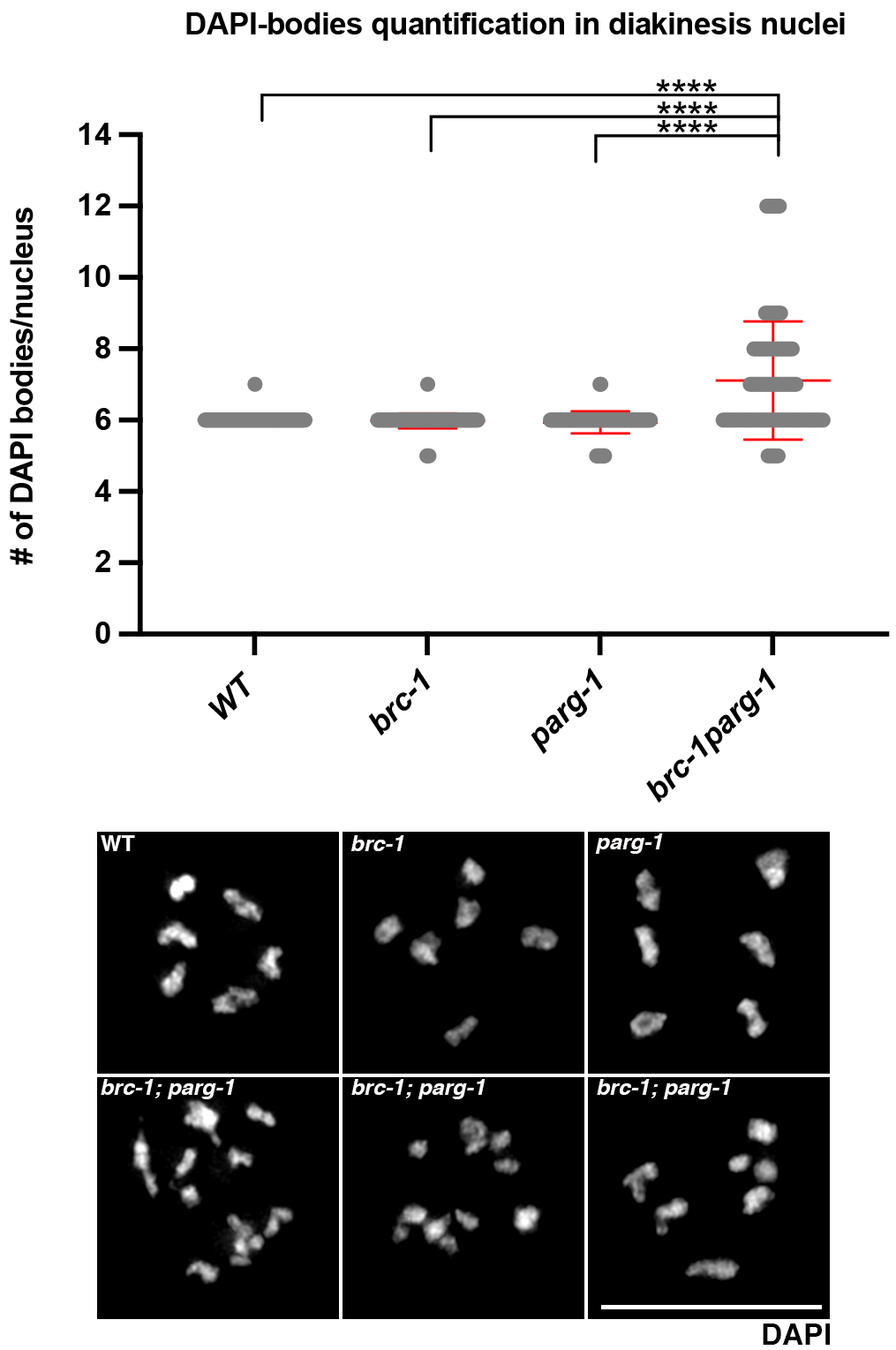

### Supplemnetal Figure 2

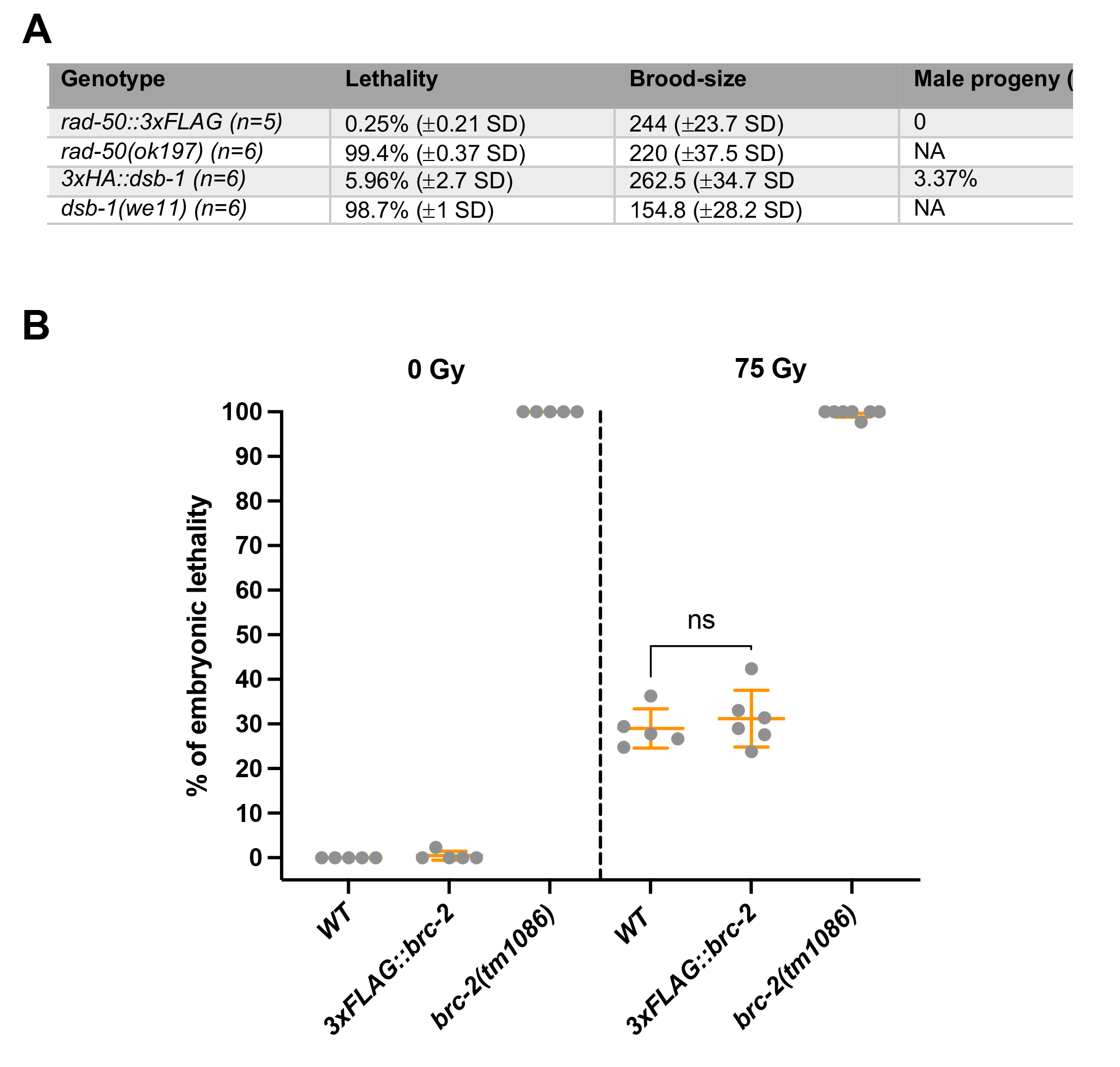

### Supplemnetal Figure 3

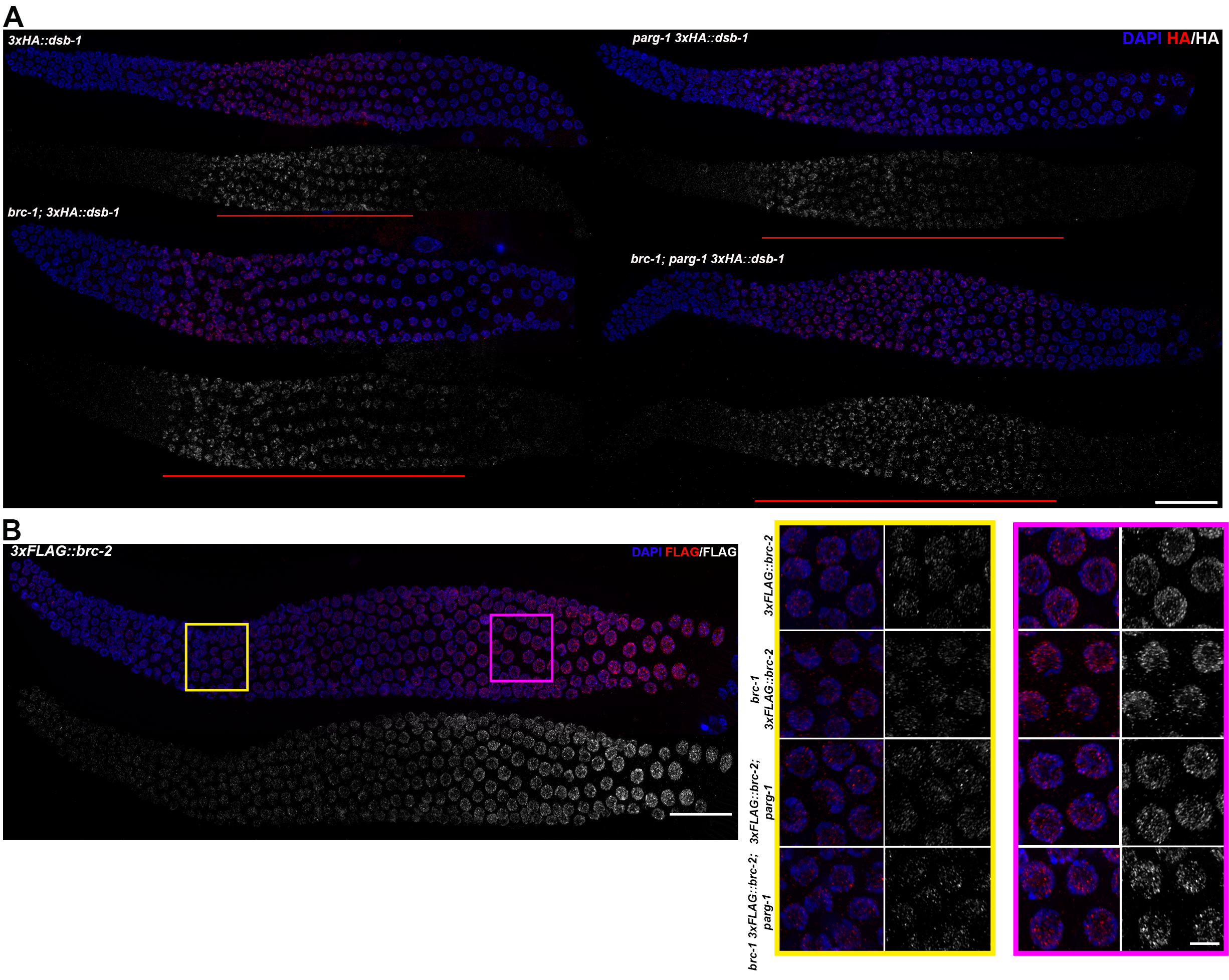

### Supplemnetal Figure 4

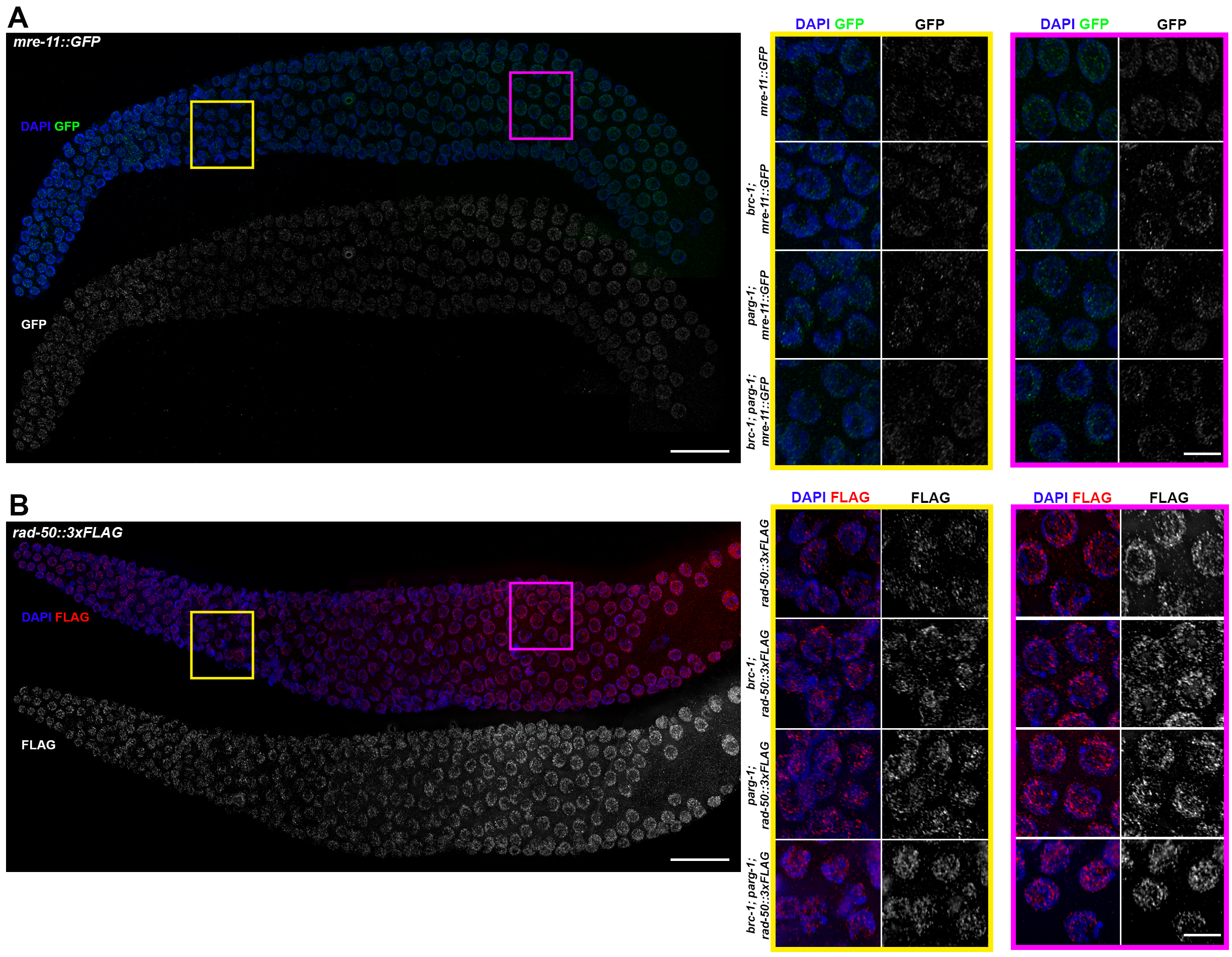

### Supplemnetal Figure 5

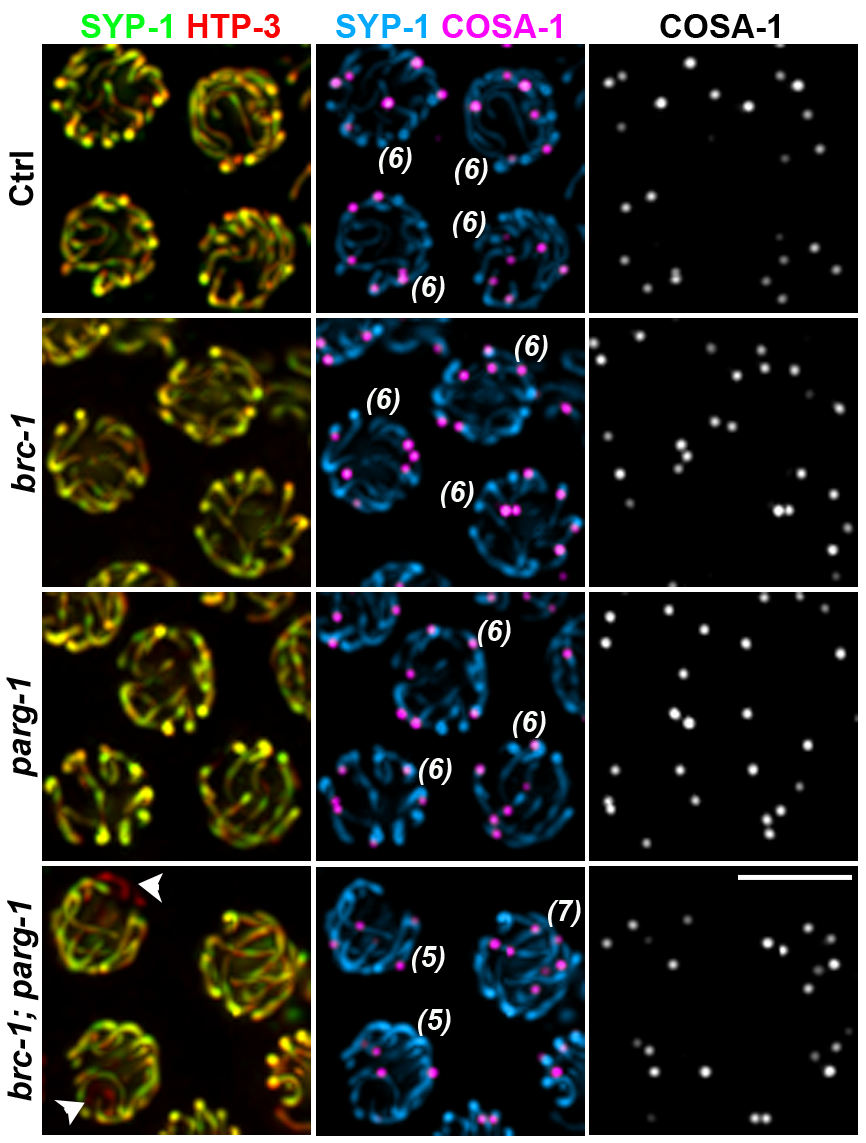

### Supplemnetal Figure 6

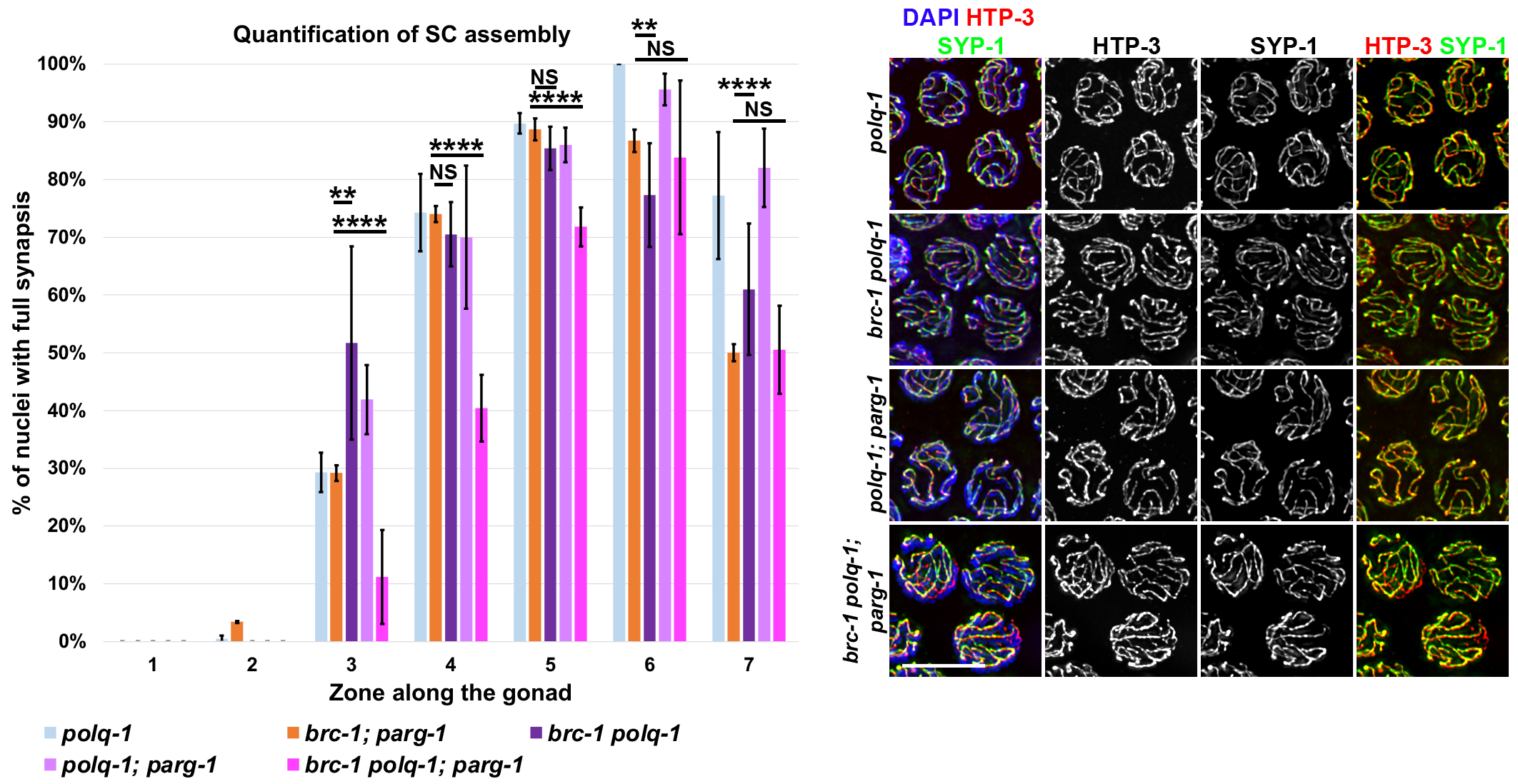
